# PerSort facilitates characterization and elimination of persister subpopulation in mycobacteria

**DOI:** 10.1101/463232

**Authors:** Vivek Srinivas, Mario L. Arrieta-Ortiz, Eliza J.R. Peterson, Nitin S. Baliga

**Affiliations:** Institute for Systems Biology, Seattle, WA, USA; Departments of Biology and Microbiology, University of Washington, Seattle, WA, USA; Molecular Engineering Program, University of Washington, Seattle, WA, USA; Lawrence Berkeley National Lab, Berkeley, CA, USA

**Keywords:** Mycobacterium, phenotypic heterogeneity, persisters, antibiotic tolerance, nutrient starvation

## Abstract

*Mycobacterium tuberculosis* (MTB) generates phenotypic diversity to persist and survive the harsh conditions encountered during infection. MTB avoids immune effectors and antibacterial killing by entering into distinct physiological states. The surviving cells, persisters, are a major barrier to the timely and relapse-free treatment of tuberculosis (TB). We present for the first time, PerSort, a method to isolate and characterize persisters in the absence of antibiotic, or other pressure. We demonstrate the value of PerSort to isolate translationally dormant cells that pre-exist in small numbers within *Mycobacterium spp*. cultures growing under optimal conditions, but which dramatically increased in proportion under stress conditions. The translationally dormant subpopulation exhibited multidrug tolerance and regrowth properties consistent with persister cells. Furthermore, PerSort enabled single-cell transcriptional profiling that provided evidence that the translationally dormant persisters were generated through a variety of mechanisms, including *vapC30, mazF*, and *relA/spoT* overexpression. Finally, we demonstrate that notwithstanding the varied mechanisms by which the persister cells were generated, they converge on a similar low oxygen metabolic state that was reversed through activation of respiration to rapidly eliminate persisters fostered under host-relevant stress conditions. We conclude that PerSort provides a new tool to study MTB persisters, enabling targeted strategies to improve and shorten the treatment of TB.

**Summary:** We have developed a novel method, PerSort, to isolate translationally dormant cells that pre-exist in small numbers within *Mycobacterium spp*. cultures growing under naïve conditions (i.e., absence of antibiotic treatment), but dramatically increase in proportion under stress conditions. The translationally dormant cells have high tolerance to isoniazid and rifampicin, and can regenerate the parental population structure in standard media, albeit after a significantly longer lag phase, indicating they are persister cells. Single-cell expression profiling demonstrated that the translationally dormant persister subpopulation is a mixture of *vapC30, mazF*, and *relA/spoT* overexpressing cells, indicating there are multiple pathways to become a persister cell. Regardless of the mechanism by which they are generated, the persister cells have reduced oxidative metabolism, which is reversed upon addition of L-cysteine to effect complete clearance by INH and RIF under host-related stress.

## Introduction

Studies have revealed cell-to-cell variability in organisms from all domains of life –unicellular to multicellular. Through cell-to-cell variability, cell populations are prepared for sudden environmental changes by harbouring subpopulations that are phenotypically pre-adapted. This evolutionary strategy, known as bet-hedging (1), confers fitness advantage to pathogens, such as *Mycobacterium tuberculosis* (MTB), which routinely experience varying environments within the host. In addition to spontaneous bet-hedging, MTB responds to host derived stresses with physiological changes (e.g., shifts in metabolism and respiration, induction of toxin-antitoxin systems, cell wall modifications) that allow it to survive and persist. These physiological changes (either stochastically- or environmentally-induced) result in antibiotic tolerance, in which MTB is genetically susceptible to antibiotics but exists in a physiological state rendering it refractory to drug killing. These persistent states are a major reason why long courses of antibiotic therapy are required to treat human tuberculosis (TB) (2); standard chemotherapy of TB requires 6 months of treatment and 5% are not cured even then (3, 4). There is an urgent need for new strategies that shorten the duration of treatment and target drug tolerant MTB. Addressing this gap requires a better understanding of how MTB generates phenotypic heterogeneity to withstand pressures from the host environment and evade antibiotic therapy.

Recent work has studied phenotypic heterogeneity in MTB from single-cell analyses using fluorescent reporter strains. Cell-to-cell variability of MTB was captured *in vitro* and during murine infections using a reporter of 16s rRNA gene expression (5). The microscopy-based platform was able to track heterogeneity in growth rate under standard growth conditions and found heterogeneity was amplified by stress conditions and murine infection. However, the non-growing subpopulation was not isolated and characterized further, most likely due to low fluorescence levels of the reporter. In another study, Jain et al developed a dual-reporter mycobacteriophage (ϕ^2^DRM) system and used fluorescent activated cell sorting (FACS) to isolate drug tolerant MTB cells from *in vitro* cultures and human sputa (6). However, the necessity to re-infect daughter cells with ϕ^2^DRM limited the ability to follow isolated cells over generations and study their regrowth patterns. The study also used a reporter that specifically enriched for persisters of isoniazid treatment (i.e., fluorescent protein fused to the *dnaK* promoter), disregarding multidrug tolerant subpopulations which are known to exist within the host environment (7).

Here, we sought to develop a fluorescent reporter system to isolate and characterize multidrug tolerant subpopulations of mycobacteria from naïve growth conditions (i.e., absence of antibiotic treatment). We wanted to avoid killing susceptible cells with antibiotics in order to study both persister cells *and* actively growing cells from the same culture and prevent drug pressure from confounding persister formation. Instead, based on a considerable body of work linking bacterial stress pathways to the acquisition of drug tolerance via translation inhibition (8-13), we developed “PerSort” to enrich for translationally dormant mycobacterium (i.e., cells that are able to transcribe but not translate a fluorescent reporter) under naïve conditions. We demonstrated the persister-like properties of the translationally dormant subpopulation and their increased abundance during stress conditions. Moreover, we performed single-cell transcriptional profiling of “PerSorted” cells and highlight various mechanisms that generate translationally dormant mycobacteria. Finally, using these mechanistic insights we revealed a physiological response common to persister cells and demonstrated that activating respiration potentiated both isoniazid and rifampicin to rapidly clear mycobacteria. Importantly, this was achieved from a host-relevant condition that fosters multidrug tolerant mycobacteria. This study confirms that PerSort enables a better understanding of mycobacterial phenotypic heterogeneity and can lead to new strategies for shortening TB treatment.

## Results

### Multidrug tolerant mycobacteria increase in nutrient starved conditions and show lag-dormancy

To establish a system that enriches for multidrug tolerant mycobacteria, we grew *M. smegmatis* (MSM) in nutrient rich (7H9 media, 0.2% glycerol, 0.05% Tween-80, and ADC complementation) and nutrient starved (PBS and 0.05% Tween-80) conditions. Deprivation of nutrients results in a marked slowing of mycobacterial growth and concurrent phenotypic tolerance to various antibiotics, including isoniazid (INH) and rifampicin (RIF) (14, 15). Moreover, studies have established the relevance of the bacterial physiological state adopted under nutrient starved conditions to TB infection (16). We incubated wild type MSM mc^2^155 strain in nutrient rich or nutrient starved conditions for 12 hours (h) then treated with either 5x minimum inhibitory concentration (MIC) INH or RIF (40 and 15 μg/mL respectively). Antibiotic-treated MSM cultures were diluted and plated over a 20 h interval. Percent survival was calculated, in reference to time 0 h, and used to generate kill curves **(Figure 1a, 1b)**. According to the bimodal model of bacterial persistence (17), multiple subpopulations were identified, either drug susceptible or drug tolerant, as distinguished by varying slopes of killing. We further estimated the abundance of drug tolerant subpopulations in either nutrient rich or nutrient starved conditions via the intersection of the y-axis with the extrapolated slope of the drug tolerant subpopulation. Thus, MSM cultures grown in standard conditions (i.e., nutrient rich) were ∼45% tolerant to INH and ∼30% tolerant to RIF. Whereas after nutrient starvation, the proportion of the drug tolerant subpopulations increased to ∼75% for INH and ∼50% for RIF. The increase in delay of RIF killing under nutrient starved conditions, might reflect cell wall alterations that are required for adaptation to nutrient deprivation, but also restrict entry of RIF (18, 19).

**Figure 1:**
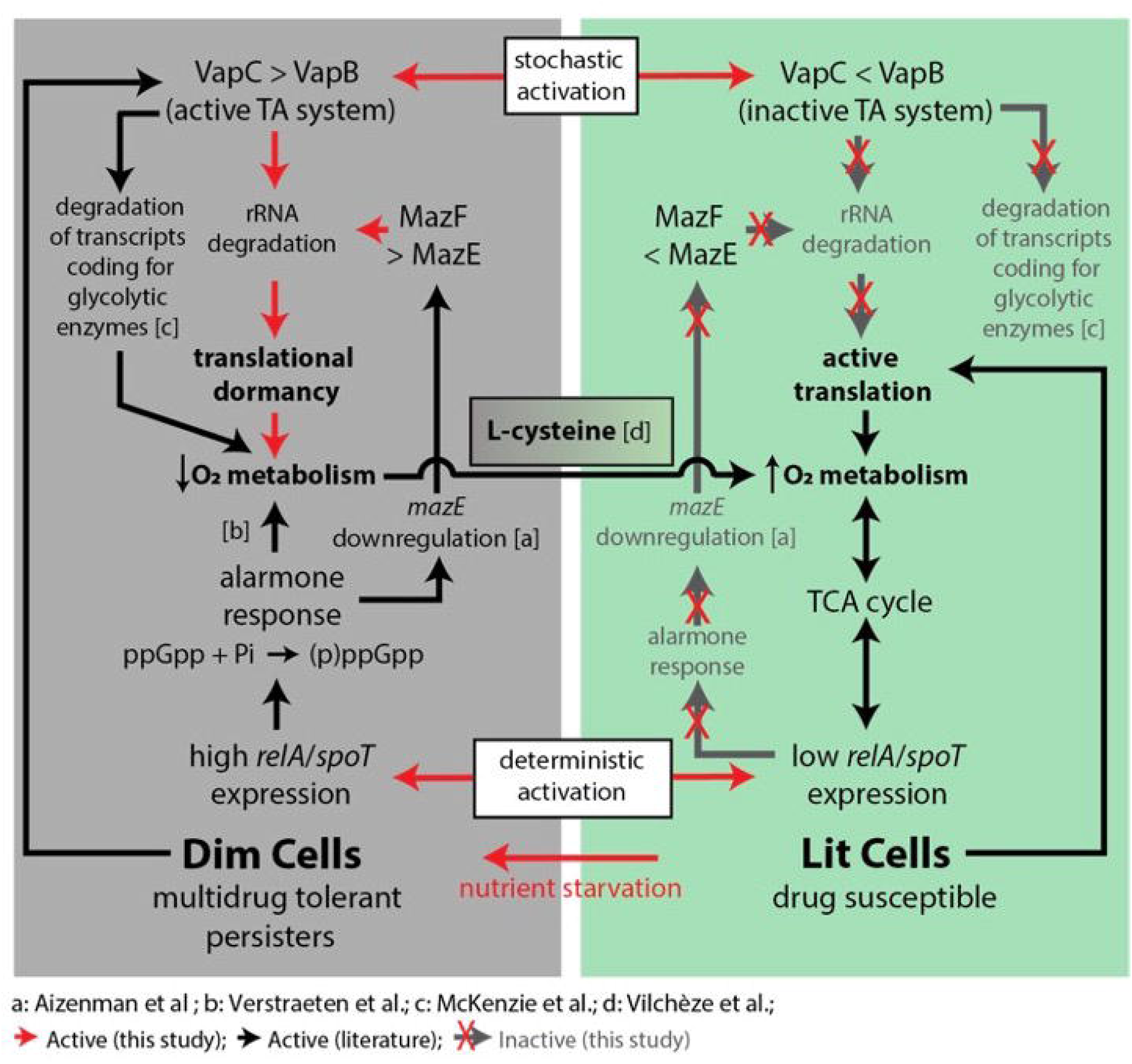
Time-kill curves and growth characteristics of *M. smegmatis* from nutrient rich and nutrient starved conditions. **a**,**b**. Time-kill curves of MSM cultures grown in nutrient rich (NR) or nutrient starved (NS) conditions treated with 5x MIC INH (**a**) or 5x MIC RIF (**b**). The solid lines indicate experimentally observed time-kill curves. Data points are from three experimental replicates, error bars were calculated by measuring standard deviation between replicates. The dashed and dotted lines distinguish the slopes of the susceptible and tolerant subpopulations, respectively. **c**. ScanLag analysis showing time of appearance (TOA) of cultures grown in nutrient rich and nutrient starved conditions. Error bars within the violin plot are standard deviation with confidence interval of 0.9. The dashed lines indicate the mean TOA from cultures grown in nutrient rich or nutrient starved conditions. Mean TOA of nutrient starved and nutrient rich conditions were compared with Students-t test.

Another advantage of the nutrient deprivation model for enriching and characterizing multidrug tolerant mycobacteria is that nutrient starved bacteria can easily grow upon being returned to nutrient rich media, thus this model allows easy quantification of drug susceptibility and growth patterns. We characterized the population wide growth characteristics of the nutrient rich and nutrient starved cultures using ScanLag, a technique that combines cell plating with high-throughput imaging (20). Following 12 h incubation in nutrient rich or nutrient starved conditions, MSM mc^2^155 cultures were plated onto nutrient rich plates (7H10 media, 0.2% glycerol, OADC) and observed for colony growth by imaging at 1 h intervals. The mean time of appearance (TOA) of individual colony forming units (CFUs) from the nutrient rich conditions was determined to be 37 h, whereas the TOA from the nutrient starved conditions was 41 h **(Figure 1c)**. This significant delay in resuming growth is due to a greater abundance of cells with lag-dormancy, a phenotype well-established with drug tolerance (21, 22), which explains why nutrient deprived mycobacteria are multidrug tolerant.

### PerSort isolates translationally dormant mycobacteria that are multidrug tolerant

To further characterize multidrug tolerant subpopulations, we developed a FACS-based protocol to isolate translationally dormant mycobacterial cells. In drug tolerance inducing conditions, translation is repressed via various mechanisms that downregulate rRNA (e.g. RelA and CarD) or degrade rRNA (e.g. VapC and other toxins) [10-15]. Therefore, we hypothesized that a fluorescent reporter that relays translation activity could efficiently sort and isolate drug tolerant mycobacterium that form in a multitude of ways. The reporter plasmid (Trans-mEos2, **Figure S1**) was constructed by inserting the highly stable fluorescent gene, *mEos2*, under the transcriptional control of an anhydrotetracycline (ATc)-inducible promoter and a strong mycobacterial translation initiation sequence (Shine-Dalgarno sequence), then inserted into the PstKi plasmid (23-25). The PstKi plasmid contains an integrase gene and thus eliminates copy number variability among cells by integrating into the mycobacterium genome. Moreover, the ATc-inducible promoter achieves detectable fluorescence from a genome-integrated reporter while also allowing for modulation of reporter expression.

The Trans-mEos2 plasmid was transformed into wild type MSM (creating the MSM-mEos2 strain) for differentiation of mEos2 fluorescence levels in mycobacterial cells using FACS. To minimize clumps, MSM-mEos2 cultures (grown in 7H9, 0.2% glycerol, 0.05% Tween-80, ADC complementation and high shaking conditions) were passed through a 10 μm filter before running on the BD FACS Influx instrument. Parameters for single-cells were defined by 3 μm rainbow fluorescent beads and the efficiency of single-cell sorting was assessed using a mixed culture of ATc-induced MSM-mEos2 cells (resistant to kanamycin) and MSM cells expressing mCherry (resistant to hygromycin) (26). Cells from the mixed fluorescent reporter cultures were sorted and plated onto 7H10 plates with appropriate antibiotic selection. Sorting was found to be 93% efficient **(Figure S2)**.

Sort gates were defined for cultivable and non-cultivable cells (i.e., dead cells and viable but non-culturable cells). Viable but non-culturable cells (VBNCs) are antibiotic tolerant, but they lack the ability to regain active growth in standard conditions and therefore do not contribute to bacteria survival (27). To exclude non-cultivable cells, MSM-mEos2 cultures were stained with propidium iodide (PI) and PI signal was used for gate selection. Heat-killed cultures were used for evidence of non-culturable cells in the PI(+) gate **(Figure S3)**. Gates for translationally active and dormant cells were defined based on mEos2 fluorescence signal in the PI(-) cultivable population **(Figure 2a)**. Wild type MSM and MSM-mEos2 cultures were grown in the presence of ATc (500 ng/mL) for 12 h and stained with 0.5% PI. Background fluorescence from wild type MSM cells was used to define the gate for translationally dormant cells (“dim”). For clear distinction between subpopulations, gates were set at a higher mEoS2 fluorescence signal to define MSM-mEos2 translationally active cells (“lit”), leaving ∼25% of PI(-) cells unselected **(Figure 2a)**. Further, the PI signal gates were set such that thresholds for selection of cultivable cells, PI(-), decreased with the mEos2 signal, as VBNCs were expected at high proportions in low mEos2 signal range. Based on these FACS gates and following ATc-induction of MSM-mEos2 cultures in standard 7H9 media (i.e., same as nutrient rich conditions) for 12 h (OD600 > 1), a majority of cells were translationally active “lit” cells, while a consistent subpopulation of translationally dormant “dim” cells was also present, reaching about 1% of the population **(Figure 2a)**. We sorted equal number of cells from the dim, lit, and PI(+) gates and plated onto 7H10 media **(Figure 2b)**. All of the cells from the dim and lit subpopulations formed CFUs, whereas only 25% formed CFUs from the PI(+) subpopulation, confirming cells from the PI(-) negative (i.e., dim and lit) gates were cultivable and devoid of VBNCs and dead cells. Finally, we measured dim cell proportions following ATc-induction of MSM-mEos2 cultures in standard growth conditions until cultures were either in exponential (∼6 h after ATc addition) or stationary (12 h after ATc addition) phase of growth. Dim cells were found in low numbers (∼0.4% of population) during exponential phase and increased during stationary phase to make up ∼1% of the population **(Figure S4)**. This indicates that dim cells pre-exist under low stress conditions (i.e., exponential phase) and a density-dependent increase in dim cell formation, likely caused by nutrient depletion of the culture during stationary phase (28).

**Figure 2:**
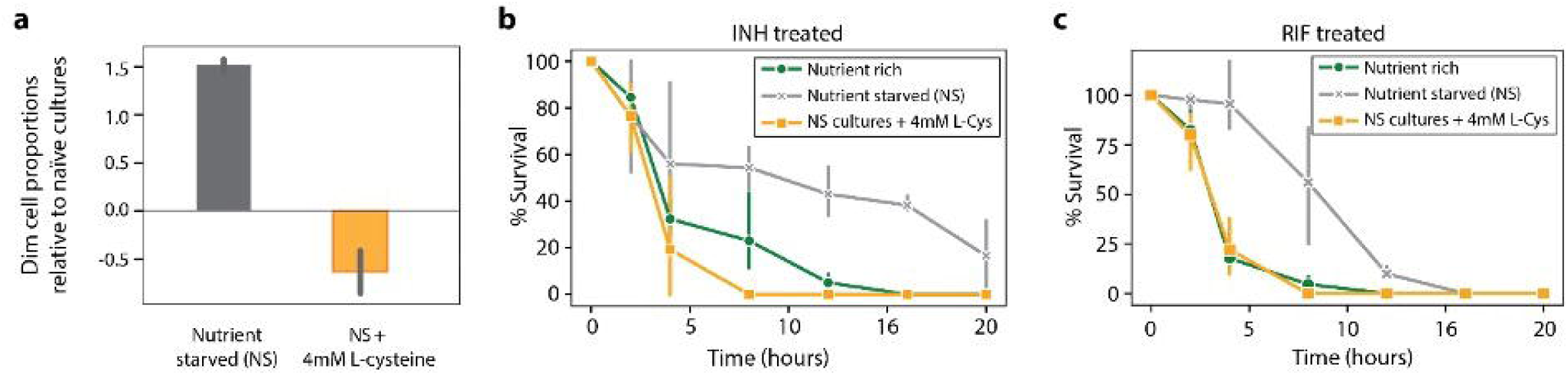
PerSort isolation and characterization of subpopulations from ATc-induced MSM-mEos2 cultures. **a**. Top: Population structure of ATc-induced (500ng/ml) MSM-mEos2 cells from single-cell gates, represented by mEos2 fluorescence on x-axis and propidium iodide (PI, 5µl/ml) fluorescence on y-axis. Polygons indicate the gates for non-culturable cells (PI(+)), dim cells, and lit cells with the proportion of cells in that particular gate. Bottom: Cumulative distribution of mEos2 florescence intensity in MSM-mEos2 cells, dashed lines indicate boundaries of sort gates for dim, lit, and unselected PI(-) cells. **b**. Cultivability of PerSorted dim, lit and PI(-) cells from ATc-induced MSM-mEos2 cultures. The “% cultivable” was calculated from the number of CFUs from 200 sorted cells. Experiments were performed in triplicates and error bars represent the standard deviation between replicates. Significance of cultivability difference between dim, lit and non-culturable subpopulations was calculated with Students t-test. **c**. ScanLag analysis of dim, unselected PI(-) and lit cells from PerSorted MSM-mEos2 cultures induced with ATc (500 ng/mL), the dashed lines in the violin plot indicate mean TOA of the sorted subpopulations. Error bars within the violin plot are standard deviation with confidence interval of 0.9. **d**,**e**. % survival of 5x MIC INH or 5x MIC RIF treatment of PerSorted dim and lit cells of MSM-mEos2 cultures induced with ATc (500 ng/ml), compared to % survival of whole populations (i.e., unsorted) of MSM-mEos2 induced with ATc (500 ng/ml) and grown in nutrient rich and starved conditions. Experiments were performed in triplicates and error bars represent the standard deviation between replicates. Significance of survival between dim and lit subpopulations was calculated with Students-t test.

Having developed the fluorescent reporter system and optimized the FACS procedure to sort translationally active and dormant cells from naïve growth conditions (i.e., absence of antibiotic treatment), we investigated the growth characteristics and drug susceptibility of the dim and lit subpopulations. For growth characterization, cells were sorted from the PI(-) gate of ATc-induced MSM-mEos2 cultures and ScanLag analysis was performed as described previously **(Figure 2c)**. The dim cells showed a longer TOA compared to lit cells, with a difference between the subpopulations (∼200 minutes) similar to that observed between nutrient rich and nutrient starved cultures (Figure 1c). As expected, cells from the unselected PI(-) gate had a mean TOA in between the dim and lit subpopulations, suggesting it is almost an equal mixture of translationally active and inactive cells. To assess drug susceptibility, 100,000 dim and lit cells from ATc-induced MSM-mEos2 cultures were sorted into 7H9 media containing either 5x MIC INH or 5x MIC RIF. The survival of dim and lit subpopulations was determined by comparing CFUs after 12 h INH or 8 h RIF treatment to CFUs before drug treatment **(Figure 2d, 2e)**. The drug treatment times were selected from the slowed phase of killing from the biphasic kill curves **(Figure 1a, 1b)**. A significantly larger percentage of dim cells survived both INH (p-value = 0.001) and RIF treatment (p-value = 0.006), indicating the dim cells are a multidrug tolerant subpopulation. Further, the survival of dim cells was compared to the entire population of ATc-induced MSM-mEos2 cultures (i.e., unsorted) grown in either nutrient rich or nutrient starved conditions. The percentage of dim cells surviving INH and RIF treatment matched closely with the nutrient starved cultures. These results reveal the striking similarities, both in terms of growth characteristics (i.e., lag-dormancy) and antibiotic susceptibility (i.e., multidrug tolerance), between dim cells and nutrient starved cultures. As such, we hypothesized the dim cells are a phenotypically heterogeneous subpopulation, with translational dormancy and multidrug tolerance, that increase in proportion under nutrient deprivation because their physiological state is best suited to withstand such environmental stress.

### Translationally dormant mycobacteria form at different probabilities in naïve versus nutrient starved conditions

We used the demonstrated capability of PerSort to isolate persister-like mycobacterial cells to investigate the population structure and regrowth characteristics of subpopulations from both naïve (i.e., absence of antibiotic treatment) and nutrient starved conditions. Following ATc induction of the MSM-mEos2 culture for 12 h, 100,000 dim and lit cells were PerSorted into standard 7H9 media. Sorted samples were grown to an OD600 of 0.6, induced with ATc again for 12 h and reanalyzed by FACS. Cultures generated from both dim and lit subpopulations had similar structure as the parent population vis-à-vis proportions of dim and lit cells **(Figure 3a)**. These results demonstrate that translational dormancy of dim cells is not heritable and dim and lit cells interconvert under standard *in vitro* growth conditions. We further assessed regrowth by measuring OD600 over time after PerSorting 100,000 dim and lit cells into standard 7H9 media. As expected, dim cell cultures grew after a longer lag phase than lit cells **(Figure 3b)**, but the subpopulations reached similar maximum growth rate and carrying capacity **(Figure 3c)**. This demonstrates that dim cells can resume normal growth (*i*.*e*., same as lit cells) in low stress conditions, which is a key characteristic of persister cells (29). Further, ATc-induced MSM-mEos2 cultures were grown in either naïve or nutrient starved conditions and then PerSorted to determine the population structure. Both dim and lit cells were present in the nutrient starved cultures, but there was a significant increase in the proportion of dim cells **(Figure 3d)**. This confirmed our hypothesis, establishing that dim cells (i.e., translationally dormant mycobacteria) dramatically increase in proportion upon nutrient starvation, further suggesting that the likelihood of generating a dim cell in each cell division is distinct under low stress versus high stress conditions **(Figure 3e)**. Whereas dim cells (and likely other phenotypically heterogeneous subpopulations) form at low probability under naïve conditions, nutrient deprivation favors the formation of dim cells (a translationally dormant and multidrug tolerant subpopulation) and enables mycobacteria to withstand stress, including drug treatment.

**Figure 3:**
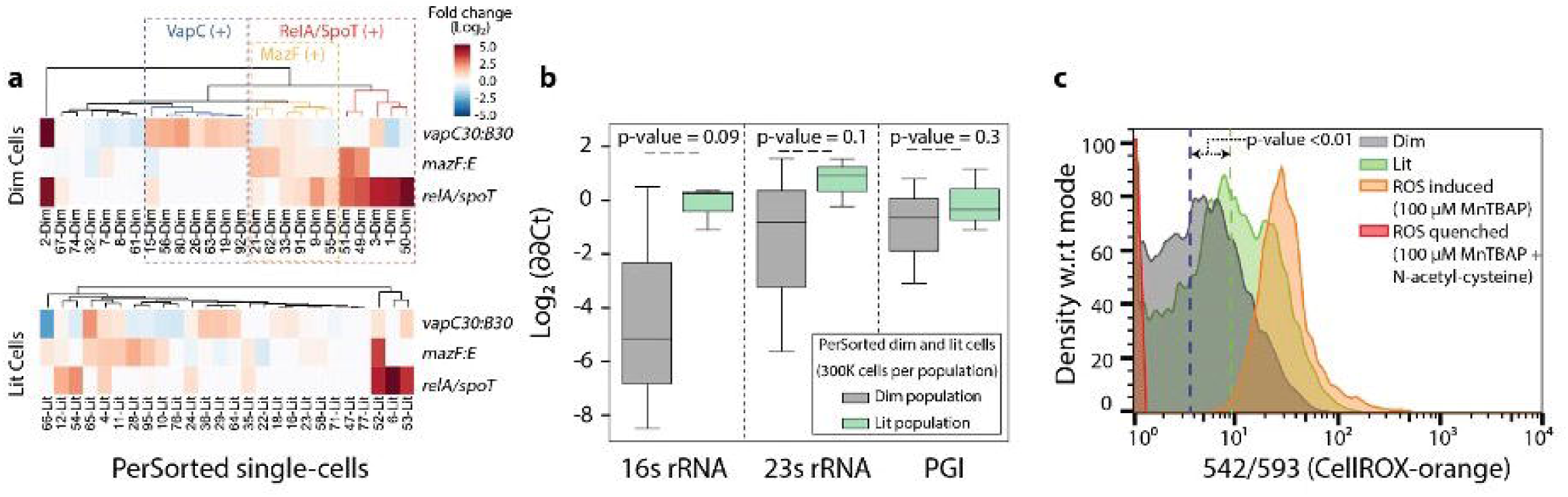
Regrowth dynamics of PerSorted dim and lit cells and population structure under naïve versus nutrient starved conditions. **a**. Dim and lit cell proportions in cultures regrown from PerSorted dim and lit populations obtained from ATc-induced MSM-mEos2 cultures. The proportions of cells in each subpopulation were measured from the gating methods described in Figure 2a. **b**,**c**. Lag phase and maximum growth rate were calculated from OD600 absorbance measurements of dim and lit cells PerSorted into 7H9 media. Error bars were calculated by measuring standard deviation in the lag phase and maximum growth rates between dim and lit cells (*n* = 100,000). Significance of differences between dim and lit subpopulations were calculated with Students-t test. NS, not significant. **d**. Population structure of ATc-induced MSM-mEos2 cultures grown in naïve (green) and nutrient starved (grey) conditions. **e**. A model describing the formation of dim cells under naïve and nutrient starved conditions in mycobacterium. The probability of dim cell formation was calculated from their relative proportion in the indicated culture conditions.

### Translationally dormant mycobacteria are composed of three discernable subtypes of *vapC30, mazF* toxins and(or) *relA/spoT* over-expressing cells

To further characterize the formation of phenotypically heterogeneous mycobacteria, we profiled within individual cells of both dim and lit subpopulations following PerSort, the transcript levels of 45 genes that were previously implicated in persister formation and drug tolerance in *Mycobacterium* spp. and *E. coli* **(Table S1)**. Single-cell gene expression profiling was performed with the Fluidigm Biomark 48×48 system per manufacturer instructions and assayed relative to single-cell genomic DNA signal. The DNA signal from single dim and lit cells was found to have low variation in expression **(Figure S5a)**, indicating that transcripts per genome copy accurately reports differential gene expression. Transcript abundances were normalized to a spike-in RNA control to account for experimental noise **(Figure S5b)**.

Kernel PCA (kPCA) with radial basis function identified distinct clusters consisting of dim and lit cells as well as some overlap between the subpopulations **(Figure S6)**. The overlap could indicate some phenotypic uncertainty, based on the expression of selected persister genes. We used a tree-based feature selection (see *Methods*) to rank the persister genes for their ability to differentiate the dim and lit subpopulations (30). Using the top features **(Table S2)**, we performed unsupervised hierarchical clustering and dynamic tree cutting to identify four distinct clusters within the dim cells **(Figure 4a)** (31, 32). Specifically, clustering revealed subtypes of translationally dormant mycobacteria with high *relA/spoT, vapC30*, or *mazF* expression (and another subtype with no distinct signature of persister gene expression). Importantly, similar clustering analysis did not detect any statistically significant subtypes within the lit cells **(Figure 4a)**. Increased expression of *relA/spoT* was observed in higher proportions of dim cells, compared to lit cells. The bifunctional activity of the protein (i.e., hydrolase and synthase) (33) could suggest different functions of RelA/SpoT between the dim and lit subpopulations.

**Figure 4:**
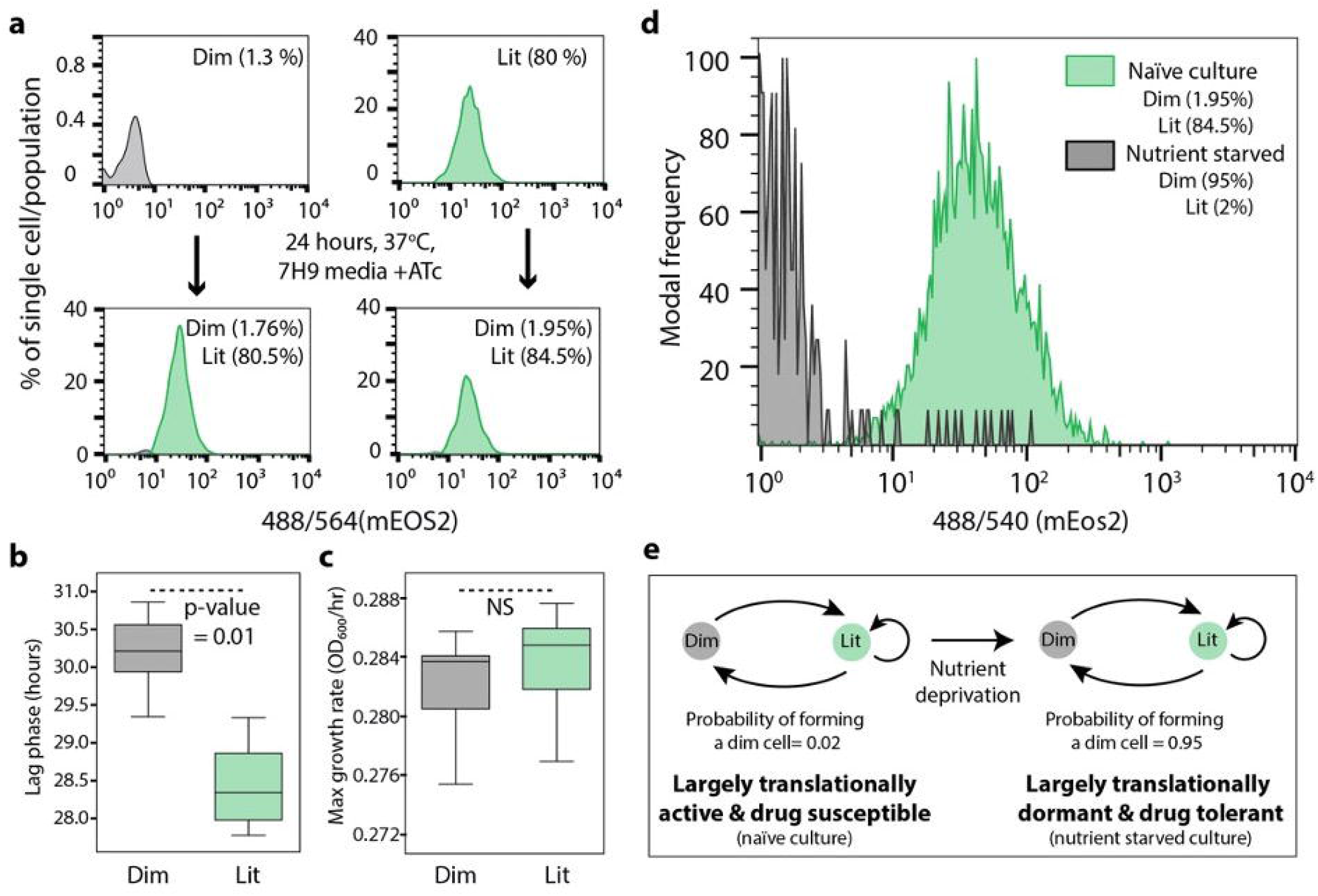
Single-cell gene expression and ROS levels from PerSorted single dim and lit cells. **a**. Hierarchical clustering of top ranked features in PerSorted dim and lit single-cells. Blocks in the cluster represent the statistically significant clusters (p-value = 0.01). **b**. qRT-PCR measurement of 16s rRNA, 23s rRNA and Phospho-glucoisomerase (PGI, MSMEG_5541) levels in PerSorted dim and lit cells (*n* = 300 cells). Error bars were calculated by measuring standard deviation of gene expression. P-values were calculated with Student’s t-test between the dim and lit cells. **c**. Reactive oxygen Species (ROS) levels, measured with CellRox, of ATc-induced MSM-mEos2 cultures grown in nutrient rich conditions and PerSorted into dim and lit subpopulations. ROS was quenched with N-acetyl cysteine and induced with MnTBAP and used as reference controls. Significance of ROS levels between dim and lib subpopulations was calculated by K-S test (K-S statistics of CellRox orange intensity between dim and lit population; K-S max Difference: 56.1%, K-S max at Intensity: 14.8551, K-S probability: >99.9%).

We sought further evidence that one or some combination of the three gene features identified from single-cell expression profiling (i.e., RelA/SpoT, VapC30, or MazF) were active in the dim cell subpopulation. The VapC30 and MazF toxins belong to a family of type II toxin-antitoxin (TA) systems, implicated in mycobacterial dormancy and persistence (28, 34, 35). Activation of the type II TA system results in toxin-mediated cleavage of tRNAs (36), or rRNAs (12, 36) causing translational dormancy (37). We analyzed 16S and 23S rRNA levels in dim and lit cells by qRT-PCR. We discovered that indeed dim cells have lower rRNA content relative to lit cells (**Figure 4b**, 16s rRNA log2 normalized FC = -4.17, 23s rRNA log2 normalized FC = -2.34). There was no discernible decrease in transcript level of a highly expressed metabolic gene (**Figure 4b**, phosphoglucoisomerase log2 FC = -0.87). These results support the notion that some of the translationally dormant mycobacteria are formed via toxin-mediated cleavage of rRNA.

### Translationally dormant mycobacteria are in a low O_2_ respiratory state

The single-cell expression data revealed that translationally dormant mycobacteria may be formed by at least three distinct mechanisms. The multiplicity of mechanisms confers robustness to the pathogen, but also thwarts therapeutic strategies to block formation of the multidrug tolerant mycobacteria. Nonetheless, we found evidence from literature that the mechanisms for persister formation converge on a common physiology and predicted the dim cells are in a reduced state of respiration. Direct measurements of respiration rate (e.g., oxygen electrode or Seahorse analysis) are problematic with low bacterial cell numbers. Therefore, to test whether dim cells have lower O_2_ metabolism we measured reactive oxygen species (ROS) levels in single-cells from the PerSorted dim and lit subpopulations using CellRox-orange. It is established that increases in ROS levels are linked with increased flux to the TCA cycle and oxidative metabolism (38, 39). Cultures of ATc-induced MSM-mEos2 were grown in standard 7H9 media, stained with CellRox-orange (40), and PerSorted to compare ROS levels between dim and lit cells **(Figure 4c)**. As reference controls, MSM-mEos2 cultures were treated with Mn-TBAP (ROS inducer) or N-acetyl-cysteine (ROS quencher) with Mn-TBAP. Cultures were incubated in CellRox-orange stain for 1.5 h to ensure uniform dye permeation across the subpopulations. The ROS levels were significantly lower in dim cells **(Figure 4c)**, confirming the translationally dormant mycobacteria have reduced oxidative metabolism.

### Activation of oxidative metabolism eliminates translationally dormant mycobacteria and achieves faster killing by INH and RIF in nutrient starved conditions

Since respiration may play an important role in generating and/or maintaining translationally dormant mycobacteria, we explored the use of L-cysteine to activate oxidative metabolism and reduce the proportion of dim cells. It is well-known that exogenous amino acids fuel the TCA cycle and oxidative respiration in bacteria, especially molecules such as cysteine and proline which are rapidly metabolized (41). Cultures of MSM-mEos2 were grown in standard 7H9 media with 4mM L-cysteine (similar growth observed in 7H9 media with or without 4 mM L-cysteine, data not shown), induced with ATc and PerSorted for dim and lit subpopulations. The L-cysteine treated MSM-mEos2 cultures were found to be completely devoid of dim cells **(Figure 5a)**. Further, we supplemented nutrient starved MSM cultures with 4 mM L-cysteine and then treated with either 5x MIC INH or 5x MIC RIF. Kill-curves for the antibiotic and L-cysteine treated cultures were generated as previously described **(Figure 5b, c)**. Activation of respiration by adding L-cysteine to nutrient starved cultures potentiated clearance by INH and RIF at rates identical to or better than rates for nutrient rich cultures. This further corroborates that, similar to dim cells, the expanded translationally dormant mycobacteria in nutrient starved conditions share a low respiratory physiologic state. Ultimately, these results demonstrate that drug adjuvants that activate respiration could shorten drug treatment, particularly in environmental niches that foster drug tolerance in mycobacteria.

**Figure 5:**
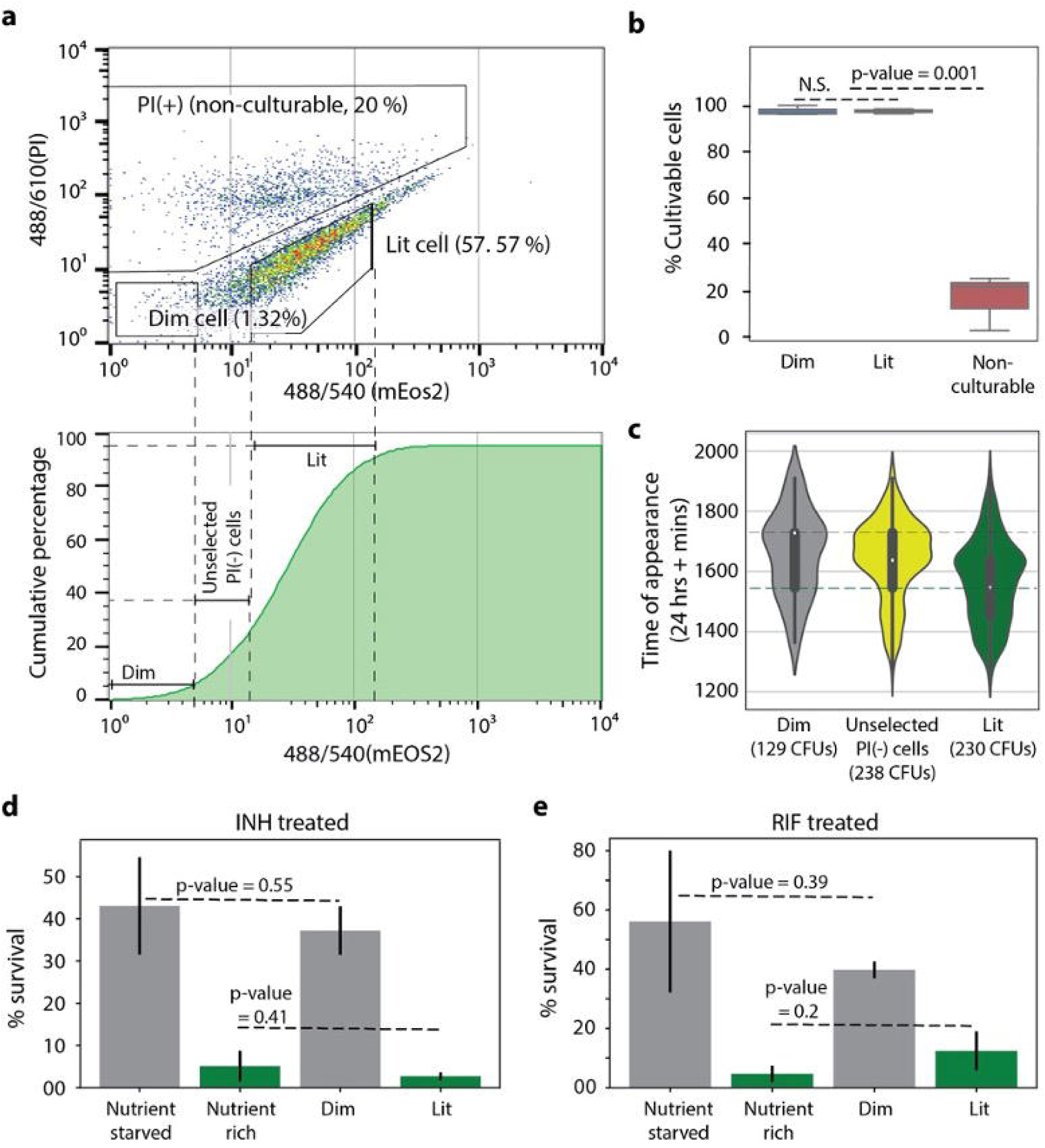
Dim cell proportion and time-kill curves upon L-cysteine addition to nutrient starved *M. smegmatis*. **a**. Proportion of dim cells in MSM-mEos2 cultures grown in nutrient starved conditions with or without the addition of 4mM L-cysteine. Dim cell proportions in each condition were calculated with respect to MSM-mEos2 cultures grown in naïve conditions. Error bars were calculated by measuring standard deviation in dim cell proportions between 10 PerSort replicates. **b**,**c** Time-kill curves of MSM cultures grown in nutrient starved conditions with 4mM L-cysteine and treated with 5x MIC INH (**b**) or 5x MIC RIF (**c**). Time-kill curve data for nutrient rich and nutrient starved conditions (no L-cysteine) is from Figure 1. Data points are averaged from three independent experiments and error bars represent the standard deviation in percent survival between the replicates.

## Discussion

The ability of microorganisms to survive sudden environmental changes stems from the formation of phenotypically heterogeneous subpopulations. Phenotypic heterogeneity confers fitness advantage to clonal microbial communities, such as infectious MTB, but impedes efforts of the immune system to clear the pathogen as well as chemotherapeutic efforts to rapidly treat TB. In this study, we developed a method to identify and characterize multidrug tolerant subpopulations of mycobacterium cells. We demonstrated that our method enables a better understanding of phenotypic heterogeneity in mycobacteria in naïve and stressed conditions, and could lead towards novel strategies to shorten TB treatment.

We generated a fluorescent reporter system, PerSort, which sorts mycobacterial cells based on translational activity. We found a small proportion of translationally dormant (“dim”) cells from naïve cultures, in the absence of stress such as antibiotic treatment. We confirmed the translationally dormant subpopulation was tolerant to both INH and RIF, demonstrated a longer lag phase upon regrowth, and could regenerate the original population structure upon regrowth (i.e., a mixture of translationally active and dormant cells). These data indicate the translationally dormant subpopulation identified by PerSort consist of cells with properties of persisters. We report these cells pre-exist in low numbers in an isogenic MSM culture growing without stress and exponential phase of growth. The translationally dormant subpopulation (along with other phenotypically heterogeneous subpopulations) is generated stochastically, as a bet-hedging strategy, for the mycobacterial population to withstand unpredictable environmental stress.

Indeed, we discovered an increase in the proportion of the translationally dormant subpopulation under density-dependent stress, and even more so under nutrient starvation. This suggests an environmentally induced component that forms this subpopulation, in addition to stochastic formation. It also begs the question of what is influencing the translational state of this mycobacterial subpopulation. The PerSort method overcomes the challenge of characterizing persisters at single cell resolution by sorting them and, importantly, not killing the susceptible cells with antibiotics. This technological advancement has overcome the confounding issue that antibiotic treatment itself induces persister cell formation (42, 43) and allows comparative analysis of persister cells *and* actively growing drug susceptible cells from the same culture. Because of these novel capabilities of PerSort, we were able to quantify transcript abundance of 45 genes associated with persister formation and drug tolerance in single-cells from both the translationally dormant (i.e., dim) and translationally active (i.e., lit) subpopulations. Expression analysis revealed that translationally dormant persisters are a mix of *vapC30, mazF*, and *relA/spoT* overexpressing cells. These results reinforce the hypothesis that there are multiple pathways (both stochastically and deterministically activated) to become a persister cell and reveals the complex and combinatorial schemes used by mycobacteria to generate heterogeneous subpopulations.

Within the translationally dormant cells, we found that high expression of toxin *mazF* was also associated with high *relA/spoT* expression. We suspect that mazEF could be regulated by the alarmone response, elicited by RelA/SpoT synthesis of (p)ppGpp, in a manner that is induced deterministically by stress conditions (44). In contrast, the single-cell expression data suggest that *vapC30* overexpression can also act to induce persister formation, in a spontaneous and alarmone-independent manner. This supports a recent study demonstrating that a *relA/spoT* knockout mutant of MSM, with reduced alarmone response, still formed persisters at levels similar to wildtype (45). This collection of evidence points toward multiple mechanisms of generating the translationally dormant mycobacteria characterized here, some of which are deterministically activated (controlled by *relA/spoT* in response to stress) and some of which are stochastically activated (controlled by spontaneous *vapC30* activation) (46). While MSM has a single VapBC-type TA system, MTB has 70 copies of VapBC (35), indicating the human pathogen has evolved to increase phenotypic diversity and bet-hedging for survival in the host environment. Much work is still needed, specifically using live cell monitoring techniques, to understand how and when these TA systems are activated to form persisters, and the contribution of other mechanisms, either stochastic or deterministic, to phenotypic heterogeneity in MTB.

The activation of TA systems (47) and the alarmone response (48) has been shown to decrease oxidative metabolism in bacterial persisters. Moreover, it is expected that translational dormancy would be associated with low respiration (28). Given this knowledge, we demonstrated lower ROS levels in the translationally dormant subpopulation compared to translationally active cells, indicating reduced oxygen metabolism in the persister cells. In other words, regardless of their mechanism of formation (i.e., *vapC30, mazF*, or *relA/spoT* overexpression), the persister subpopulation shares a low oxygen respiratory state, which presents a vulnerability that could be targeted to modulate this subpopulation **(Figure 6)**. We confirmed that addition of L-cysteine, which is known to activate oxidative metabolism (49), dramatically reduced the proportion of translationally dormant cells in nutrient starved MSM cultures, conditions where the population is abundant. This goes beyond previous studies by directly demonstrating that promoting oxidative metabolism reduces the proportion of multidrug tolerant persister cells that form independent of drug pressure. Furthermore, the addition of L-cysteine was able to effect complete clearance by INH and RIF in nutrient starved conditions. The addition of L-cysteine potentiates faster drug killing by converting (or limiting the formation of) multidrug tolerant persister cells, cells with enhanced fitness advantage in the host-relevant stress (nutrient starved) conditions, to a population with increased oxidative metabolism and drug susceptibility. Our findings prove that until properties of heterogeneous subpopulations are disrupted, we will not be able to successfully clear mycobacterial cells in infected patients. This study highlights how novel methods to isolate and characterize heterogeneous subpopulations can enable targeted strategies to eliminate detrimental persister cells, and thereby shorten the course of treatment.

**Figure 6.**
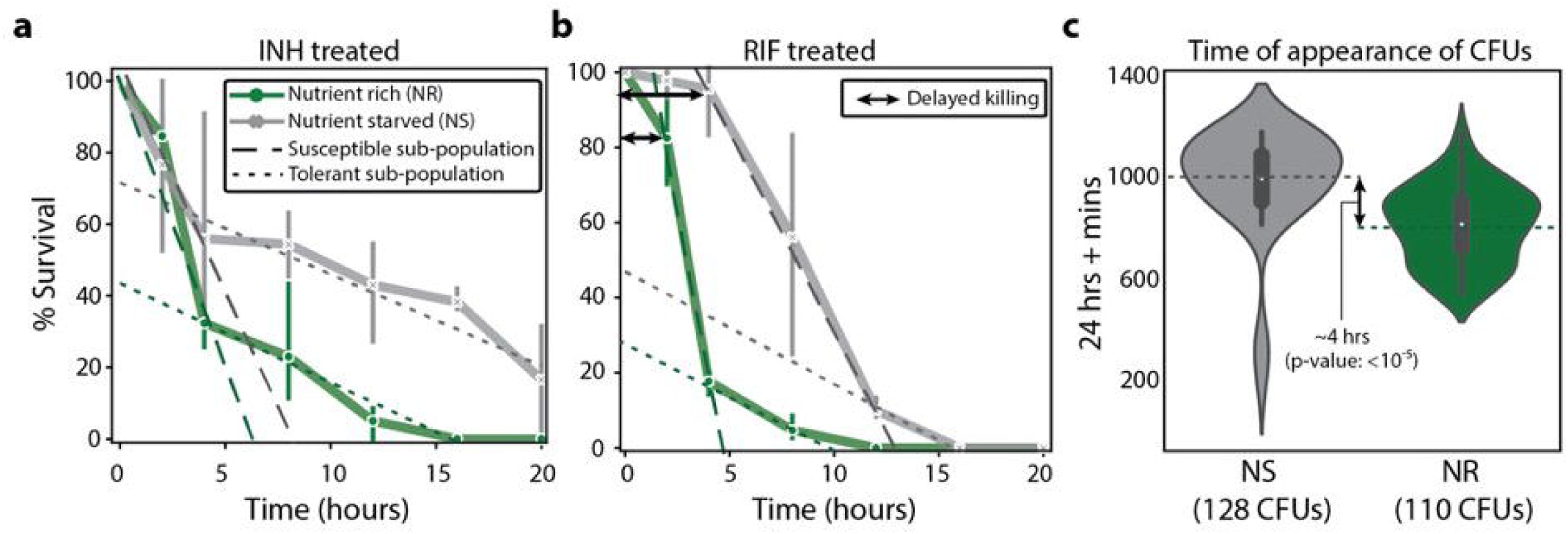
The persister subpopulation converges on a low O_2_ metabolic state, which was exploited by addition of L-cysteine to convert the subpopulation to drug susceptible cells. The diagram depicts the processes driving dim (multidrug tolerant persisters) and lit (drug susceptible) cell formation, as discovered through single-cell expression analysis. Evidence from literature further connects these various mechanisms to the O_2_ respiratory state of either dim or lit cells. Red arrows indicate connections identified in this study, black arrows are previously determined connections from literature (references indicated), and white boxes are proposed mechanisms for persister cell formation.

## Materials and Methods

### Bacterial growth and MSM-mEos2 strain development

*Mycobacterium smegmatis* mc^2^ 155 strain (MSM) was obtained from ATCC and grown in 7H9 broth medium (Difco) with 0.2% glycerol, 0.05% Tween-80, and 10% ADC enrichment (BD biosciences). pSTKi-mEos2 plasmid was constructed from pSTKi plasmid and pRSETa mEos2 plasmid (23, 25). Synthetic oligo with translation initiation signal (mycoSD) was used to amplify mEos2 from pRSETa mEos2 plasmid, and amplified fragment was inserted into the pSTKi plasmid with restriction ligation at BamH1 and EcoR1 sites. The pSTKi-mEos2 transcript was electroporated into electrocompetent MSM cultures, transformed colonies were selected on 7H10 plates with 30 µg/ml kanamycin (KAN). MSM-mEos2 cells were cultured in 7H9 media supplemented with 30 µg/ml KAN.

### Development of PerSort

The PerSort method was developed using BD FACS Influx. A 70 micron tip and sheath fluid from BD bioscience was used in sorting and FACS analysis. Green fluorescent beads (3.5 micron) and 5 micron Accudrop beads from BD biosciences were used to calibrate the instrument for laser alignment, compensation, and cell sorting. Propidium Iodide (PI) stain (SigmaAldridge, 0.5%) was used to stain dead cells. Heat killed cells (incubated at 70° – 80° C for 5 mins) were used as dead cell control for PI stain. MSM cells with pCHERRY3 plasmid (MSM-mCherry) (26) was used to optimize single-cell mycobacterium sorting. MSM-mEos2 (KAN resistant) cultures were grown to OD600 of 0.6 and induced with 500 ng/ml ATc. Induced cultures were incubated at 37° C for 12 h. MSM-mCherry (Hygromycin resistant) were mixed with induced MSM-mEos2 cultures in equal proportions and single mEos2 and mCherry cells were sorted and plated on 7H10 plates with Kan and Hyg to determine the efficiency of single-cell sorting **(Figure S2)**. Gates for sorting dim and lit cells of ATc induced MSM-mEos2 strains (induced for 12 h) were determined using uninduced MSM-mEos2 strains stained with PI. FACS data was analyzed in FlowJo software (ver. 10).

### Antimicrobial tolerance assay

Minimum inhibitory concentrations (MIC) of the MSM-mEOS2 strains were determined with disk diffusion assay (50). To determine percent survival, cultures (bulk and PerSorted) were incubated in 7H9 media containing desired MIC concentrations of INH and RIF. Drug treated bacteria were washed with or diluted (1:100) in 7H9 media and plated (100 µl) or spotted (5 µl) on 7H10 media with 30 µg/ml KAN at time points 0, 8, 12, 16, 20, and 24 h after incubation. Percentage survival was calculated with respect to 0 h.

### qRT-PCR of PerSorted samples

Dim and lit cells were sorted into 500 µl TriZol and RNA was extracted with DirectZol RNA purification kit (Zymo Research) with the manufacturer’s instructions. qRTPCR was performed with primers mentioned in **Table S3** with Luna® Universal Probe One-Step RT-qPCR Kit (New England Biolabs). Log2 normalized RNA abundances were calculated using ValidPrime signal as reference (51).

### Single-cell persister gene expression and analysis

The Fluidigm Biomark system with 48×48 plates was used for this study. Single dim and lit cells were PerSorted into 96 well plates with VILO™ reaction mix (5x), SUPERase (Invitrogen™), and 10% NP40 in a pre-noted random order to avoid sampling bias. Sorted plates were spun down and freeze-thawed 3 times on dry ice to rupture cells. ValidPrime assay (51) for non-transcribed genomic DNA was used to determine the rupture efficiency and calculate the signal from a single nucleic-acid strand (used as reference to calculate transcript abundance). Reverse transcription (RT) was performed on freeze thawed cells with VILO cDNA preparation mixture, T4-Gene32 protein, and random hexamer primers. RNA spike-in (ECC2_SpikeIn RNA, 10 pM) was included in the RT master mix. cDNA of the genes of interest **(Table S1)** was pre-amplified with TaqMan® PreAmp master mix (Invitrogen™) and equimolar mixture of forward and reverse strand primers designed for the genes of interest **(Table S3)**. Primers were removed with Exonuclease I (Invitrogen™). Primers sets used for preamplification were primed into the 48×48 Biomark assay plates. Quantitative PCR assay with Biomark prescribed protocol was run on diluted, pre-amplified Exo 1 treated cDNA with Sso Fast EvaGreen Supermix (Bio-Rad laboratories). Quality control for determining, sorting, cell lysis, and cDNA preparation was performed by comparing the CT values of genomic DNA control and spike-in control in all the cells **(Figure S5a**,**b)**. Expression levels of genes was measured as ∂CT in individual cells with reference to genomic DNA control (expected to result from 1 copy of genomic DNA), less than or equal expression (of genomic DNA control) was considered as zero expression and assays with CT >40 were flagged as missing values. ∂CT values for each cells were corrected by adding or subtracting the deviation from median ∂CT of spike-in control for a particular cell. Hierarchical clustering was performed with Python Seaborn package and the R package, PVclust.

### Feature selection of single-cell clusters

Ratio of toxin expression to their respective antitoxin expression was calculated and included along with the other persister gene expression values. Key features that differentiate dim and lit subpopulations were selected with tree-based feature selection tool (30). Weights of the features that signify its ability to differentiate between dim and lit sub populations were estimated as an average over 100 iterations of randomly selected subsets of the dataset (70% of gene expression values). Constraints of gene regulation was also used in selection of features as most persisters specific responses are a part of alarmone response, a broad, genome wide change in transcript levels (52). The three features with the highest weights and evidence of additional criteria were selected for identification of clusters within dim and lit cells.

### In situ ROS level measurements with CellRox assay

MSM-mEos2 cells were induced with ATc (500 ng/ml) and incubated for 12 h under standard growth conditions. 10 µM N-acetyl cysteine (NAC), an oxidative agent that reduces level of in situ reactive oxygen species (ROS) was added to the induced MSM-mEos2 cultures and incubated at 37° C for 90 mins to prepare the negative control. Along with negative control, another set of induced cultures were treated with a ROS inducing 100µM MnTBAP (Sigma Aldrich, 55266-18-7) and incubated for 15 mins to prepare positive control. CellRox orange (Thermo Fisher, C10443), was added to the induced samples and control and incubated for 30 mins at room temperature. FACS analysis was performed to measure the CellROX orange intensity. ROS + and negative gates were determined with control samples and ROS levels in dim and lit cell population were determined with reference to the controls.

### Dim cell proportion and antimicrobial tolerance following L-cysteine addition

MSM-mEos2 cells were grown in PBS and 0.05% Tween-80, with and without 4mM L-cysteine, to an OD600 of 0.6. Cultures were induced with ATc (500 ng/ml) and incubated for 12 h. Dim cell proportions in both conditions were measured with PerSort assay and normalized to the dim cell proportions observed in optimal growth conditions (growth in 7H9 media without L-cysteine). Tolerance of cultures grown in various conditions was measured with time kill curve assays, under 5x MIC RIF (17.5 µg/ml) and 5x MIC INH (20 µg/ml) treatment. The % survival was measured at varying time point by counting number of CFUs on 7H10 plates in reference to the time point 0 of the respective samples.

## Acknowledgements

We thank members of the Baliga lab for critical discussions; Amardeep Kaur for her technical expertise; Tim Petersen and Monica Orellana for help with FACS. Funding was provided by the National Institute of Allergy and Infectious Diseases of the National Institutes of Health [R01AI128215] and [U19AI10676, U19AI135976]; and National Science Foundation [1518261, 1565166, and 1616955].

## Author contributions

V.S. designed research, performed experiments, analyzed data and wrote the paper. M.L.A-O. performed computational analyses. E.J.R.P and N.S.B. designed research, analyzed data, and wrote the paper.

