## Supplementary document for "PerSort facilitates characterization and elimination of persister subpopulation in mycobacteria"

**This file includes:**

Figures S1 to S6

Table S1-S3

Supplementary References

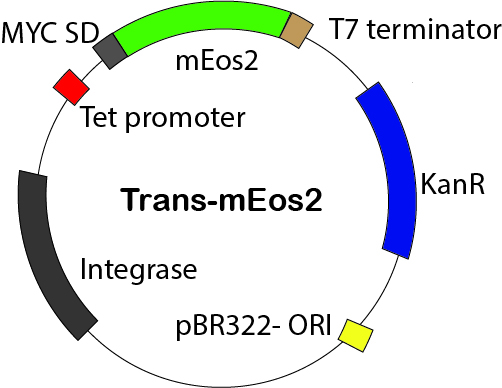

**Figure S1**. Plasmid map of Trans-mEos2. Tet promoter, tetracycline inducible promoter; MYC SD, Shine-Dalgarno sequence of *Mycobacterium spp*.; ORI, origin of replication; and KanR kanamycin resistance cassette.

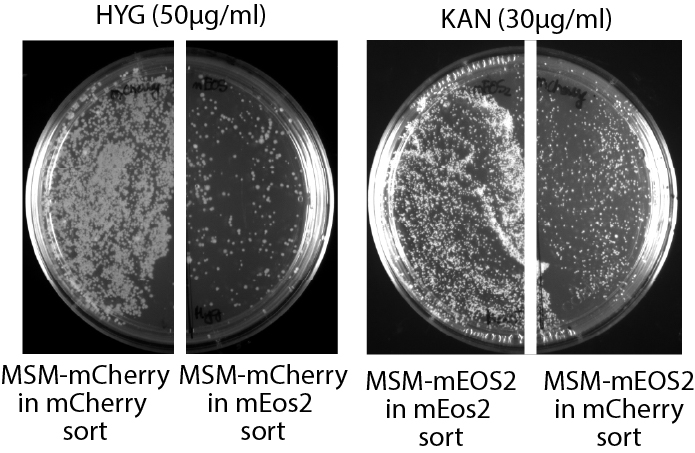

**Figure S2**. Efficiency in bacterial single-cell sorting. Bacterial cells (*n* = 2000) from the equal-proportional mixture of MSM-mCherry strain (with mCherry fluorescence and hygromycin (HYG) resistance) and MSM-mEos2 strain (with mEos2 fluorescence and kanamycin (KAN) resistance) were sorted based on mCherry and mEos2 fluorescence. Cells (*n* = 2000) from the mCherry sort and mEos2 sorts were plated onto both 7H10+HYG and 7H10+KAN plates to determine the percentage of MSM-mEos2 cells in mCherry sort and MSM-mCherry cells in mEos2 sort (i.e., inappropriately sorted). Approximately 120 colony forming units (CFUs) of MSM-mCherry were enumerated in the mEos2 sort (HYG plates, right panel) and 150 MSM-mEos2 cells in mCherry sort (KAN plates, right panel). Suggesting that only ~7% of cells were inappropriately sorted and the sorting efficiency for single bacterial cells to be ~93%.

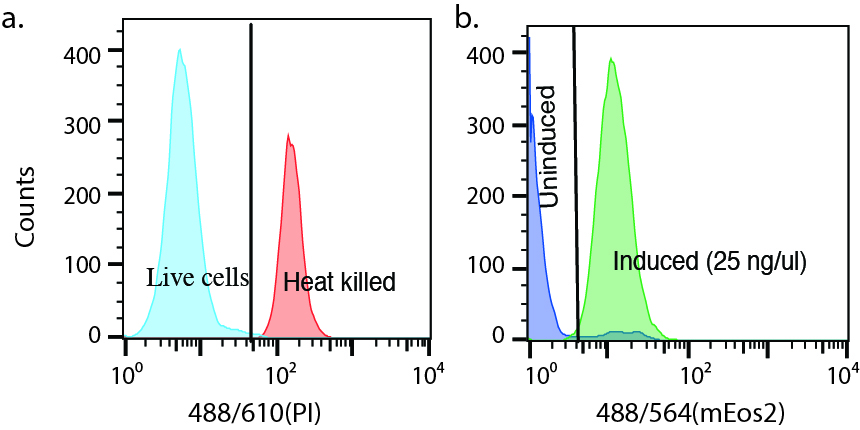

**Figure S3.** (**a**) Gating strategies used in identification of translationally dormant (dim) cells. Gating between PI stained live and dead MSM-mEos2 strains (left panel), and PI stained non-fluorescing uninduced and ATc (500 ng/ml) induced MSM-mEos2 strains (right panel). (**b**) mEos2 reporter expression in Per-Sorted MSM-mEos2 dim and translationally active (lit) sub-population.

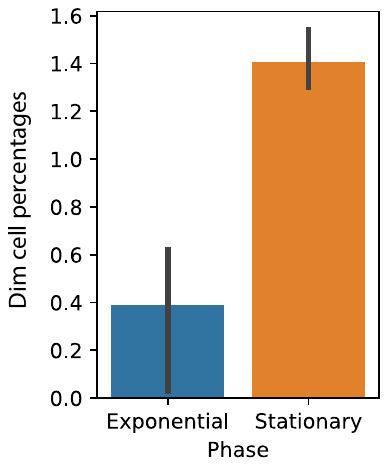

**Figure S4**. Translationally dormant (dim) cell percentages in exponential and stationary phase cultures of MSM-mEos2 cultures induced with 500 ng/ml ATc at OD600 of 0.05 and 0.6, respectively. Both samples were induced with ATc for 12 hours at 37^o^ C before they were PerSorted for dim cell percentages. Experiments were performed in triplicates, error bars were calculated by measuring standard deviation in dim cell percentages from the three replicates.

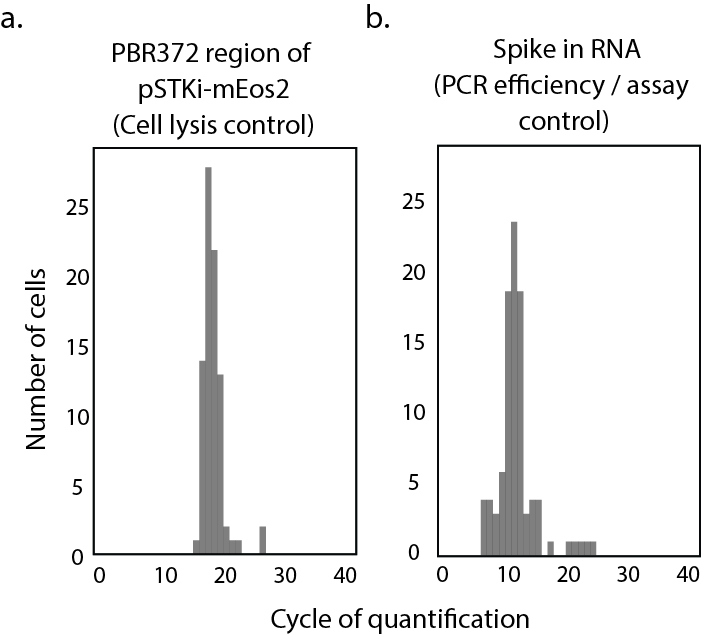

**Figure S5**. Quality analysis of Fluidigm Biomark persister gene assay. Cycle of quantification (CT) histograms for **a**. Genomic DNA expression in PerSorted dim and lit cells. Genomic region considered for measurement was from PBR372 region of pSTKi-mEos2 plasmid that is inserted into the genomic DNA of MSM-mEos2 strain. **b.** RNA spike in control from PerSorted dim and lit cells indicating uniformity in cell lysis and PCR efficiency, respectively.

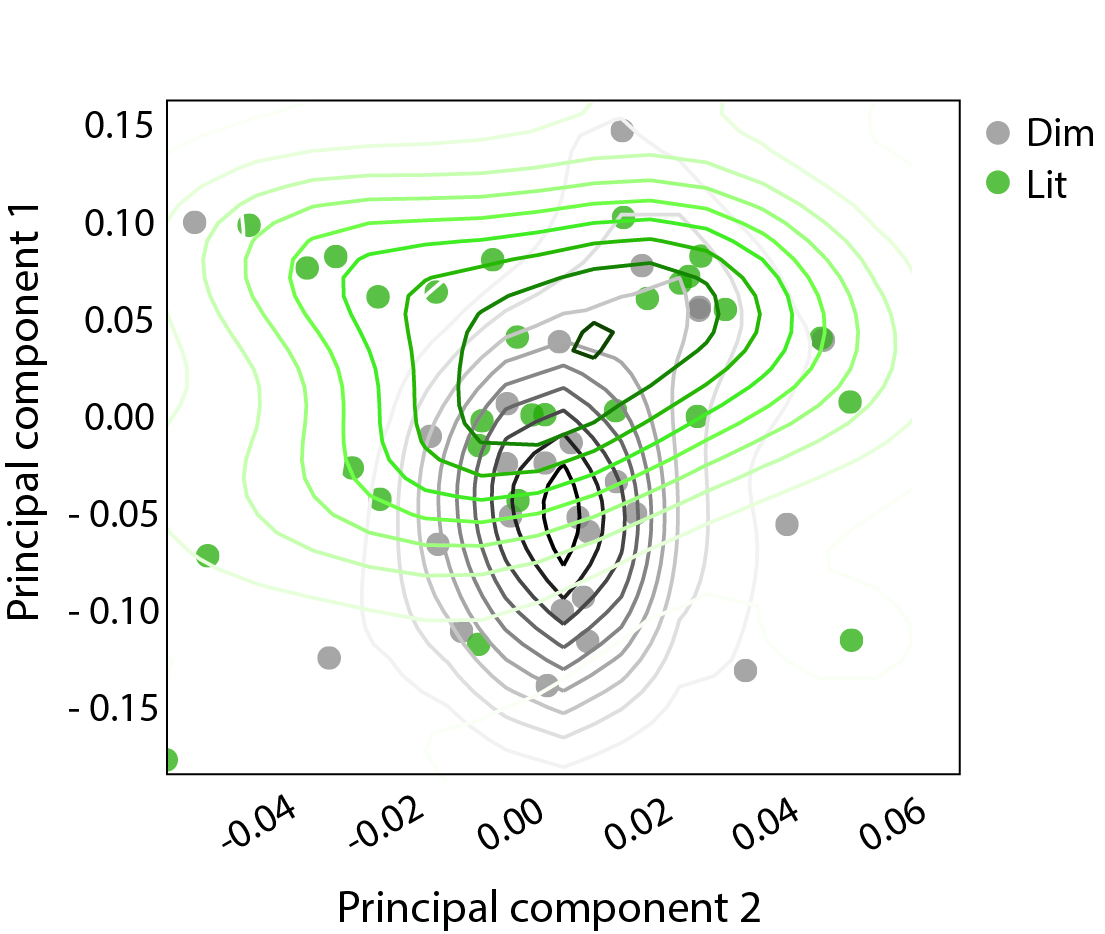

**Figure S6.** kPCA plot that used radial basis function for dimensionality reduction of persister gene expression (∂∂Ct; 45 genes) in Per-Sorted single dim and lit cells isolated from MSM-mEos2 cultures induced with ATc (500 ng/ml). Solid lines in the scatter plot indicate the density estimates of the population (outliers were omitted in the plot)

**Table S1**. Genes used in single cell persister gene expression assay

| **MSM gene ID** | **Gene name** | **Gene function** | **References** |
| --- | --- | --- | --- |
| *MSMEG_2817* | ABC efflux pump 1 | Alarmone response / membrane depolarization | (1,2) |
| *MSMEG_5659* | ABC transporter1 | Alarmone response / membrane depolarization | (1,2) |
| *MSMEG_5660* | ABC transporter2 | Alarmone response / membrane depolarization | (1,2) |
| *MSMEG_3945* | ABC transporter3 | Alarmone response / membrane depolarization | (1,2) |
| *MSEMG_2740* | *lexA* | Alarmone response | (1,2) |
| *MSMEG_2965* | *relA/spoT* | Effector of alarmone response | (1–3) |
| *MSMEG_2391* | *ppk1* | Alarmone response | (1,2) |
| *MSMEG_1633* | *dnaE2* | DNA replication | (1,2,4) |
| *MSMEG_3151* | *inhA* | Drug target | (1,2) |
| *MSMEG_2723* | *recA* | Drug target | (1,2) |
| *MSMEG_4427* | efflux pump 1 | Efflux pump 1 | (1,2) |
| *MSMEG_3944* | *devR c1* | Metabolic control | (1,2) |
| *MSMEG_6384* | *katG* | Metabolic control | (1,2) |
| *MSMEG_5244* | *devR c2* | Metabolic control | (1,2) |
| *MSMEG_1605* | *phoY1* | Persister phenotype associated gene | (1,2) |
| *MSMEG_5776* | *phoY2* | Persister phenotype associated gene | (1,2) |
| *MSMEG_1915* | *rshA* | Persister phenotype associated gene | (1,2) |
| *MSMEG_4265* | *lamA/mmpS3* | Persister phenotype associated gene | (1,2) |
| *MSMEG_0913* | *umaA* | Persister phenotype associated gene | (1,2) |
| *MSMEG_5773* | *desA1* | Persister phenotype associated gene | (1,2) |
| *MSMEG_0916* | *desR* | Persister phenotype associated gene | (1,2) |
| *MSMEG_5248* | *desA2* | Persister phenotype associated gene | (1,2) |
| *MSMEG_0424* | *hsp* | Persister phenotype associated gene | (1,2) |
| *MSMEG_4466* | *uspA* | Persister phenotype associated gene | (1,2) |
| *MSMEG_5141* | *narK2* | Persister phenotype associated gene | (1,2) |
| *MSMEG_1803* | *rsbW* | Persister phenotype associated gene | (1,2) |
| *MSMEG_6225* | proton antiporter efflux pump | Alarmone response / membrane depolarization | (1,2) |
| *MSMEG_2389* | *mdp1* | Regulator | (1,2) |
| *MSMEG_0709* | *dnaK* | Regulator | (1,2) |
| *MSMEG_2752* | *sigB* | Sigma factor | (1,2,5) |
| *MSMEG_1914* | *rpoE1* | Sigma factor | (1,2,5) |
| *MSMEG_2758* | *mysA* | Sigma factor | (1,2,5) |
| *MSMEG_0713* | *hspR* | Sigma factor | (1,2,5) |
| *MSMEG_1277* | un-annotated | TA operon (Antitoxin) | (1,2) |
| *MSMEG_1283* | *vapB30* (Antitoxin) | TA operon (Antitoxin) | (1,2,6,7) |
| *MSMEG_3180* | antitoxin | TA operon (Antitoxin) | (1,2) |
| *MSMEG_4175* | *arsR* (Antitoxin) 1 | TA operon (Antitoxin) | (1,2) |
| *MSMEG_4447* | *aze* (Antitoxin) | TA operon (Antitoxin) | (1,2) |
| *MSMEG_6762* | *arsR* (Antitoxin) 2 | TA operon (Antitoxin) | (1,2) |
| *MSMEG_1278* | un-annotated | TA operon (Toxin) | (1,2) |
| *MSMEG_1284* | *vapC30* (Toxin) | TA operon (Toxin) | (1,2,6,7) |
| *MSMEG_3181* | Toxin | TA operon (Toxin) | (1,2) |
| *MSMEG_4176* | *arsR* (Toxin) 1 | TA operon (Toxin) | (1,2) |
| *MSMEG_4448* | *mazF* (Toxin) | TA operon (Toxin) | (1,2) |
| *MSMEG_6760* | *arsR* (Toxin) 2 | TA operon (Toxin) | (1,2) |

**Table S2: Ranking of features to distinguish presorted dim and lit population of MSM-mEos2**

| **Gene name** | **Weights**  **(ability to sort dim and lit cells)** | **Evidence for gene regulation** |
| --- | --- | --- |
| *relA/spoT* | 0.07176533 | Yes |
| *vapC30* (Toxin) | 0.06485776 | Yes |
| *mazF:E* | 0.03787879 | Yes |
| *vapC30:B30* | 0.03710575 | Yes |
| *vapB30* (Antitoxin) | 0.02164502 | Yes |
| *lexA* | 0.02164502 | Yes |
| *dnaK* | 0.02164502 | Yes |
| toxin | 0.02142857 | Yes |
| *arsR* (Antitoxin) 2 | 0.01739332 | Yes |
| *mazF* (Toxin) | 0.01731602 | Yes |
| *sigB* | 0.01670431 | Yes |
| *arsR* (Antitoxin) 1 | 0.01298701 | Yes |
| ABC transporter1 | 0.00629673 | Yes |
| *ppk1* | 0.004329 | Yes |
| Proton antiporter efflux pump | 0.09848485 | No |
| ABC transporter2 | 0.06648114 | No |
| *desA1* | 0.06095327 | No |
| *hsp* | 0.05773218 | No |
| *rsbW* | 0.05764885 | No |
| *devR c1* | 0.04743867 | No |
| *hspR* | 0.04312949 | No |
| *mdp1* | 0.03690567 | No |
| *mazE* (Antitoxin) | 0.02497503 | No |
| *lamA/mmpS3* | 0.02345779 | No |
| ABC efflux pump 1 | 0.02164502 | No |
| Efflux pump 1 | 0.02071066 | No |
| Unknown | 0.01636905 | No |
| *desA2* | 0.01628403 | No |
| *dnaE2* | 0.01623377 | No |
| *mysA* | 0.01082251 | No |
| *umaA* | 0.00773037 | No |
| *inhA* | 0 | No |
| *phoY2* | 0 | No |
| *rshA* | 0 | No |
| *uspA* | 0 | No |

**Table S3**. Primers used in the study

| **MSM gene ID** | **FP** | **RP** |
| --- | --- | --- |
| *MSMEG_1277* | GGCGGATGACCTGTCGCTGA | GGCGACCCTGCGCTTGTG |
| *MSMEG_1278* | CCATGCGCTGGTCGACGGTA | TCCTCCGCGCGATCGACATC |
| *MSMEG_1283* | GAGGCGGTGGTGATGGCACT | GAGGCGGTGGTGATGGCACT |
| *MSMEG_1284* | GCGGTGGCTGACGATCCTGT | GAGTTCGCGACCACCTGGCT |
| *MSMEG_3180* | TCGCGGTGCTCATGGACGAC | CGGGCACGTGCACTGCATTC |
| *MSMEG_3181* | CCGGACGCGGTCTACGTGTT | AAGCTCACCCGCACGATCCC |
| *MSMEG_4175* | TGCCCTGGTCGACGGTGAAC | GACCTCGCGCAGCACCTTGA |
| *MSMEG_4176* | GGCACGGTGCTTCGCTTCAC | CGGTCGAAGAAGGCGTGGGT |
| *MSMEG_4447* | ACCGAGTACGCCGACATCGC | GGCGGCGACCAACTCAGACT |
| *MSMEG_4448* | AATCGAGCCAACGCCAGCCA | CGACACGGTGCGTCAAGCTG |
| *MSMEG_6760* | CCCGGACGGCGAGAAGTACG | AGCGAACCCGTCGAGGAACG |
| *MSMEG_6762* | CACGAGGCGCGACATCATGC | AGCAGGCCGGCTTTCTCCAG |
| *MSMEG_2694* | CACTTTCGCGCGTGCTCACC | GCCATGGACACCTGCGGGAT |
| *MSMEG_4671* | GATCGCCCGCAAGAGCGAGA | GACGCCTGCGTACCCTCCAG |
| *MSMEG_4466* | CGGTTCCGCCACATCACCCT | GGTGAGCGCGTAGACGGTGT |
| *MSMEG_1803* | GTGGATCCCGGTCCCGATGC | CAGGCCCGGACCCATCTGTG |
| *MSMEG_1633* | AGTGGGCCCGCATGGAGAAC | CCGAGGCCCAGCATGTCGAA |
| *MSMEG_6384* | CCGGTGAGCGTGACCTGGAG | TGCGGATCCGGATTGCCGTT |
| *MSMEG_2389* | CACAGAAGCTCCCGGCCGAT | CACAGAAGCTCCCGGCCGAT |
| *MSMEG_3151* | TCGACGGTGTGGTGCACTCG | GCGCGTCGAAGAACGGGTTG |
| *MSMEG_2723* | CAGGCGCTGCGCAAGATGAC | CTCGGGCGAGCCGAACATCA |
| *MSMEG_2740* | GACACCGGCGAGTTCACGGA | TCGAGGATGGTGCGCTGACG |
| *MSMEG_2965* | GTGCTCGCCGACGAGAAGGT | GTGCTTCGGGTCGCCCATCT |
| *MSMEG_1915* | GCCTGCGGCATTACGGCATC | TCGTGGTGCGGCTGATCTGG |
| *MSMEG_4265* | GCCGACGTGGCGCTCTATGA | GCCGACGTGGCGCTCTATGA |
| *MSMEG_5659* | TGGCGTCTCGGCCTGATGTG | TACGTGCGGGCGGATTCGTT |
| *MSMEG_0913* | GGAAGTCCGCCTGCAGGGTT | GGGTAGCGCTCGGCCTTGAA |
| *MSMEG_5660* | GTCGACCATCCGCCGGTTCA | CAGCAGGATCGCGGTGACCA |
| *MSMEG_0709* | GCGACCTCCGGTGACAACCA | GATGCCCGAGCTGCCCTTGA |
| *MSMEG_0424* | AATCCGACGGCCGCACCTAC | CCGGCGACCCGTACCTTCAG |
| *MSMEG_0713* | ACCTGCTGCGAGAGGTGCAG | AGCGCGTCGACCTGATTGGT |
| *MSMEG_3932* | CATGCGCTCGGTGACACTGC | TCTCCACCGCGACACGCTTC |
| *MSMEG_5141* | GGTCGGATCGTTGGGACGCA | CAGGATCGACGCGACCGTCA |
| *MSMEG_3945* | ATCCACGGCGAGTCGAAGGC | CGTCGAGAGGCTGCGTCGAA |
| *MSMEG_0880* | GGTCGGCAACGAGGGTGTCA | CTCGGCGTCGGTCACGAAGT |
| *MSMEG_1583* | CATCGCCGATCGCGTCAAGC | AGCCGCTCCTGCAGCTTCTC |
| *MSMEG_2391* | GCTGTTGGAGCGCGCGAAAT | AGCGCACCGACAGACCCATC |
| *MSMEG_3944* | CGACCCGAAGTCGCGGTTCT | GTGGCCTCGTCGGACGTGAA |
| *MSMEG_5244* | TCACCCAGCAGGAGCGTGTG | CGCGCCGCGATCTGTTTGTT |
| *MSMEG_1605* | CCGATCTGACGCTGGCCGAA | GTGCATGGACCCGACGACCA |
| *MSMEG_5776* | CACGCGGGATCCGGAGAAGG | CCACTCGCGGTCCATCAGCA |
| *MSMEG_2752* | TCGACATGCCGGTCGGAACC | GGCGGACATGGCCTCGGAAT |
| *MSMEG_1914* | GTCCAACGCCGAGCACTCCT | GCTTGCAACGCGGCCTTGAT |
| *MSMEG_2758* | AGGGCGAGAAGCTGCCAGTG | GCAGGTTCGCCTCCAGCAGA |
| *MSMEG_2817* | TGGGAGCCGCTGGCTTCTAC | CCGACGACGGTACCGAGGAA |
| *MSMEG_4427* | GCGGTTTGGCTTCCGCAGTC | GCGGCGCTGACCTTCAACAC |
| *MSMEG_6225* | GGGTGCCGTGGTGTCGATGA | CCAGGCCCGTGACCATCAGG |
| *MSMEG_5773* | GGCCTCGACATCGCGCCGAA | GCGGAGCACCGGCATCACGA |
| *MSMEG_0916* | CCGTCGGCGGTGTCCAGCTC | ATGCCGCACAGCGCGTACCA |
| *MSMEG_5248* | GCGGGCCTCGACGTGATCGG | CTCGGCGACGTTGGCGACCT |
| Assay control (Spike in) | TCCAGATTACTTCCATTTCCGC | GCTGGATGCCGACGCCCGTAT |
| Genomic DNA control or Valid prime control (PBR372 region of pSTKi mEos2) | TGGCTGCTGCCAGTGGCGAT | GCCCGACCGCTGCGCCTTAT |
| *MSMEG_3757* (16s rRNA) | AGAGTTTGATCCTGGCTC | GCCATGCGACCAGCAG |
| *MSMEG_3756* (23s rRNA) | GGATGCCTTGGCACTG | AGACGCCTATATATTCAGC |
